# The phosphorylation of Pak1 by Erk1/2 for cell migration requires Arl4D acting as a scaffolding protein

**DOI:** 10.1101/2024.12.02.626348

**Authors:** Ting-Wei Chang, Ming-Chieh Lin, Chia-Jung Yu, Fang-Jen S. Lee

**Author notes:** Contributed to the manuscript equally (Ting-Wei Chang, Ming-Chieh Lin). To whom correspondence should be addressed: Prof. Fang-Jen S. Lee.

## Abstract

Activation of extracellular signal-regulated kinases 1 and 2 (Erk1/2) at the plasma membrane typically results in their translocation to other intracellular sites for substrate targeting. This targeting requires scaffolding proteins to bring Erk1/2 and their substrates together. In the case of platelet-derived growth factor (PDGF)-induced signaling that activates Erk1/2 to phosphorylate Pak1 for cell migration, Erk1/2 remain at the plasma membrane. Thus, it has been unclear in this case whether a scaffolding protein is needed for substrate targeting by Erk1/2. The small GTPase Arf-like protein (Arl) 4D also promotes cell migration by targeting Pak1 to the plasma membrane. However, as this recruitment also results in the phosphorylation of Pak1, it has been unclear how this phosphorylation is achieved. Furthermore, because both Erk1/2 and Arl4D promote the role of Pak1 in cell migration, the relationship between these two processes has also been unclear. Addressing these outstanding questions, we show that Arl4D acts as a scaffolding protein by recruiting Erk1/2 and Pak1 to the plasma membrane for their assembly into a protein complex. Our findings identify Arl4D as a novel regulator of Erk1/2, reveal a conserved role for scaffolding proteins in substrate targeting by Erk1/2, as well as uncovering a previously unknown interplay among Arl4D, Erk1/2, and Pak1. Moreover, as all these factors are known to be involved in cell migration, we have shed new mechanistic insights into how this fundamental cellular process occurs.

## Introduction

Extracellular signal-regulated kinase (Erk) participates in multiple fundamental cellular processes that include cell proliferation, cell survival and cell migration (Guo *et al*, 2020; Mendoza *et al*, 2015; Ramos, 2008; Sharma *et al*, 2003). Erk exists in two major isoforms, Erk1 and Erk2. How Erk1/2 act has been studied extensively in the context of intracellular signaling through the mitogen-activated protein kinase (MAPK) cascade (Buscà *et al*, 2016; Hommes *et al*, 2003; Pearson *et al*, 2001). This process is initiated by growth factors binding to their receptors on the cell surface. Receptor activation results in the Ras small GTPase being recruited to the plasma membrane, which then recruits the Ras effector Raf, followed by MAPK kinase (MEK) 1/2 and then Erk1/2 (De Luca *et al*, 2012; Katz *et al*, 2007; Mor & Philips, 2006). This recruitment of Erk1/2 leads to their activation, for which one extensively studied fate involves Erk1/2 translocating from the plasma membrane to the nucleus in targeting their substrates (Mebratu & Tesfaigzi, 2009; Ramos, 2008). Scaffolding proteins participate in this targeting by promoting the interaction of Erk1/2 with their substrates. Activated Erk1/2 can also translocate to other intracellular sites that include the cytoskeleton, the mitochondria, and the Golgi. Scaffolding proteins have also been found to promote the targeting of Erk1/2 to their substrates at these sites (Wainstein & Seger, 2016; Yao & Seger, 2009).

Another general fate of activated Erk1/2 involves them staying at the plasma membrane. The platelet-derived growth factor (PDGF) activates Erk1/2 through the MAPK cascade (Yoshimura *et al*, 2005). Erk1/2 then targets Pak1 for phosphorylation at the plasma membrane (Sundberg-Smith *et al*, 2005b). Pak1, being also a kinase, is then activated to target its substrates in promoting cell migration (Yuan *et al*, 2010). However, whereas Erk1/2 has been shown to phosphorylate Pak1 at its T212 residue, a direct connection between this phosphorylation and the role of Pak1 in promoting cell migration remains to be better established. It has also been unclear whether a scaffolding protein is needed for the targeting of Pak1 by Erk1/2.

The Arl4 (ADP ribosylation factor-like 4) small GTPases, which is comprised of Arl4A, Arl4C, and Arl4D, act in multiple cellular processes that include cell migration (Chen *et al*, 2020a; Donaldson & Jackson, 2011; Harada *et al*, 2021; Li *et al*, 2007). Arl4A and Arl4D (Arl4A/D) have been found to promote cell migration by regulating the recruitment of Pak1 to the plasma membrane (Chen *et al*., 2020a). This Arl4A/D-mediated recruitment of Pak1 also results in the Pak phosphorylation, but the identity of the kinase that acts in this phosphorylation process has remained elusive. It is also unclear whether a relationship exists between how Arl4A/D activate Pak1 versus how Erk1/2 activate Pak1. We address these outstanding questions in the current study.

## Results

### Erk1/2 are novel interacting proteins of Arl4A/D on membrane

We initially sought to identify novel proteins that interact with Arl4A/D at the plasma membrane. To mimic the membrane-bound form of Arl proteins, we pursued a general approach of attaching proteins to liposomes. This involves tagging the protein with an 6xHis epitope and incorporating a nickel-labeled lipid into liposomes, which upon incubation leads to the protein becoming attached to liposomes. Moreover, by appending the epitope tag to the N-terminus of Arl4 proteins, we mimicked how they are oriented to the membrane through the insertion of their N-terminal myristate. We then incubated the liposome-bound Arl4 proteins with cell lysates, followed by mass spectrometry to identify novel interacting proteins. Furthermore, to select those that bind specifically to the activated form of Arl4D, we compared proteins pulled-down using liposomes bound with Arl4D-Q80L (GTP-locked form) versus those bound with Arl4D T35N (nucleotide binding-deficient form) (Fig. 1A and Dataset. EV1).

**Fig. 1.**
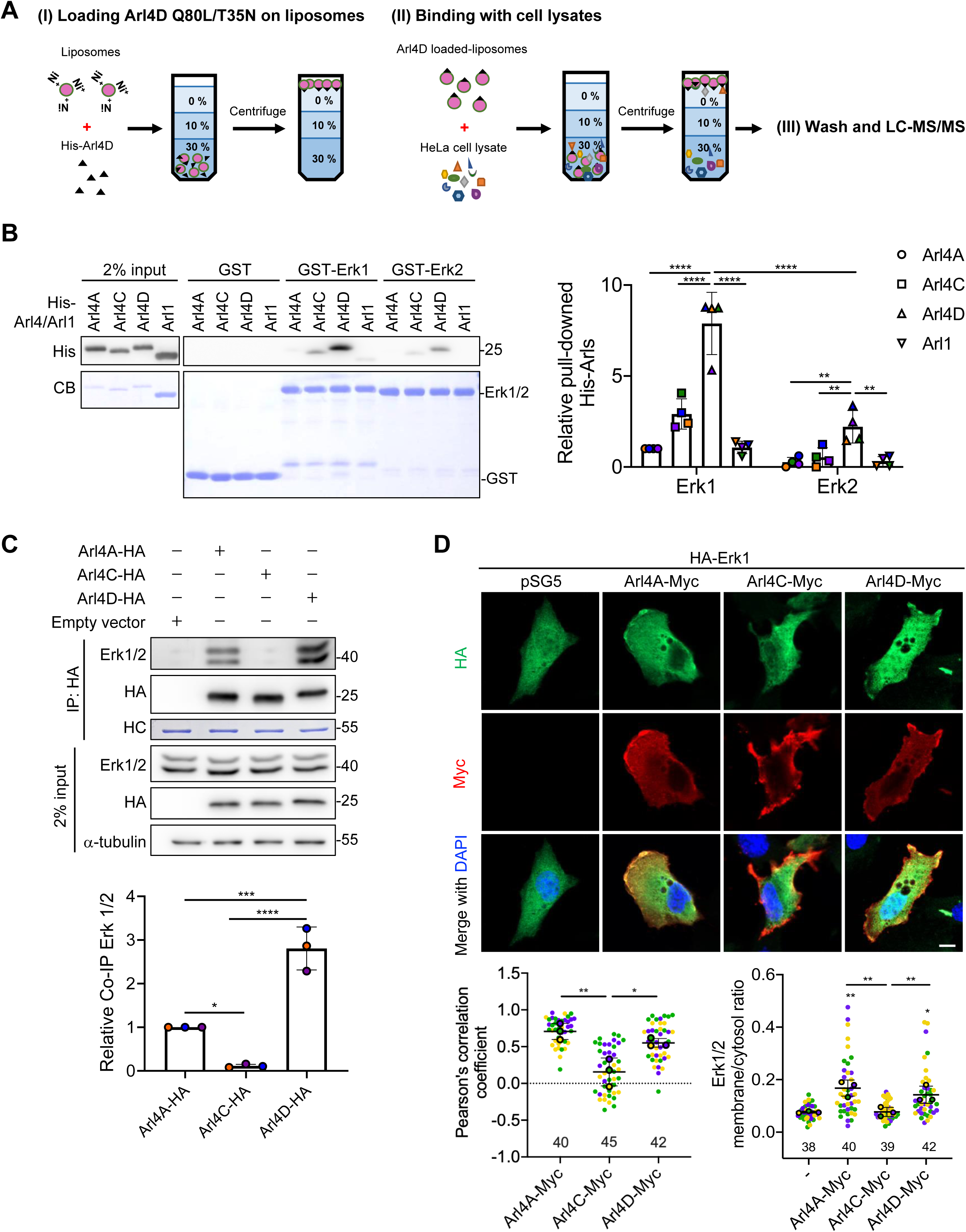
Erk1/2 is a novel and direct binding partner of Arl4D in the membrane compartment. (A) 50 mg of recombinant His-Arl4D Q80L (GTP-bound form) or Arl4D T35N (nucleotide-free form) purified from E. coli were loaded onto 0.4 mg of NTA-incorporated liposomes and driven through 30%, 10% and 0% sucrose gradient (I). The top layer containing liposomes loaded with Arl4D was mixed with 50 mg HeLa cell lysate and re-suspended to separate the Arl4D-specific interacting proteins (II). Liposomes containing Arl4D and Arl4D membrane-interacting proteins were washed through the sucrose gradient and subjected to LC/MSMS (III). (B) In vitro binding of His-tagged Arl4s and Arl1 with GST, GST-Erk1 or Erk2. His-Arl4s and Arl1 proteins pulled down from GST fusion proteins were analyzed by Western blotting. Equal amounts of GST proteins were detected by staining with CB (Coomassie blue). The His signals of the pulled-down His-Arl4s and Arl1 proteins were quantified in the dot plots, with error bars indicating the mean ± (n=4). **P < 0.01; ****P < 0.0001 (two-way ANOVA with Tukey’s post-hoc multiple comparison test). (C) NIH3T3 cells transfected with empty vector pSG5 or Arl4s-HA were subjected to co-immunoprecipitation (Co-IP). The Co-IP signals of endogenous Erk1/2 proteins were normalized with Arl4s-HA proteins after subtracting the background signal of the pSG5 control group, and the quantified results are shown in the dot plots with error bars indicating the mean ± SD (n=3). *P<0.05; ***P<0.001, ****P < 0.0001 (one-way ANOVA with Tukey’s post hoc multiple comparison test). HC, heavy chain of the antibody. (D) NIH3T3 cells transfected with the indicated proteins were stained with anti-Myc (red) and anti-HA (green) antibodies and DAPI (blue; stains the nuclei). Scale bar, 10 mm. Pearson correlation coefficients of Arl4s-Myc and HA-Erk1 signals were calculated using ZEN imaging software, respectively, and the results were shown in the dot plots with error bars indicating the mean ± SD (n=3, cells analyzed in each biological replicate were marked in the same color, and the total number of cells is indicated in the graph). The ratios between plasma membrane and cytosol of HA-Erk1 in each group were quantified as described in the Materials and Methods, and the results were shown in the dot plots with error bars indicating the mean ± SD (n=3, the cell samples are the same as described). *P<0.05; **P<0.01 (one-way ANOVA with Tukey’s post hoc multiple comparison test).

One protein identified by this approach was Erk2, which was of particular interest, as it is known to phosphorylate Pak1 at its T212 residue (Chen *et al*., 2020a; Sundberg-Smith *et al*, 2005a). Because Erk1 and Erk2 share 84% sequence similarity, we purified both and then assessed whether they interact directly with purified Arl4 proteins. *In vitro* binding assays showed that Arl4D had the strongest interaction with Erk1/2 among the Arl4 members tested, with Arl4D binding better to Erk1 than to Erk2. (Fig. 1B). We confirmed these associations by performing co-precipitation studies on cell lysates, finding that Arl4D also had the strongest interaction with Erk1/2 (Fig. 1C). Moreover, Arl4D bound preferentially to Erk1 over Erk2, when the levels of Erk1/2 in cell lysates were normalized. We also performed microscopy and found that Erk1 had better colocalization with Arl4A/D at the plasma membrane than with Arl4C (Fig. 1D).

### Erk1/2-dependent phosphorylation of the T212 residue in Pak1 requires Arl4D

As PDGF stimulation results in the phosphorylation of the T212 residue in Pak1 (Kingsley *et al*, 2002; Sundberg-Smith *et al*., 2005a), we next examined whether the Arl4D-Erk1 interaction underlies how this occurs. Erk1/2 activity is often dysregulated in cancer cell lines, with PDGF stimulation resulting in variable activation of Erk1/2. Thus, we initially sought to identify a cell line that shows Erk1/2 activity to respond robustly to PDGF stimulation. Assessing for Erk1/2 activation through their phosphorylation (p-Erk1/2), we identified NIH3T3 and A-10 cells to be such cell lines, which was further confirmed by treating these cells with U0126, a pharmacologic inhibitor of MEK1/2 (which phosphorylates Erk1/2), as this treatment led to both Erk1/2 activation and the T212 phosphorylation of Pak1 to be markedly reduced (Fig. EV1A). Thus, we focused on these cells for subsequent studies. Performing siRNA treatments, we found that PDGF-triggered phosphorylation of the Pak1 T212 residue was more efficiently reduced by targeting against Arl4D than Arl4A (Fige. 2A and EV1B). To test whether Erk1/2 require the membrane localization of Arl4D to activate Pak1, we compared the effect of expressing Arl4D wild-type (WT) versus that of a mutant (G2A) which cannot localize to membranes due to defective myristoylation. Performing siRNA against Arl4D followed by rescue with either form of Arl4D, we found that the WT, but not the G2A mutant, was able to restore the phosphorylation of the Pak1 T212 residue (Fig. 2B). Consistent with these results, overexpression of Arl4D WT, but not G2A, further enhanced Erk1/2-dependent T212 phosphorylation of Pak1 upon PDGF stimulation (Fig. 2C). Thus, these results suggested that membrane-bound Arl4D is required for Erk1/2 to phosphorylate Pak1 at its T212 residue upon PDGF stimulation.

**Fig. 2.**
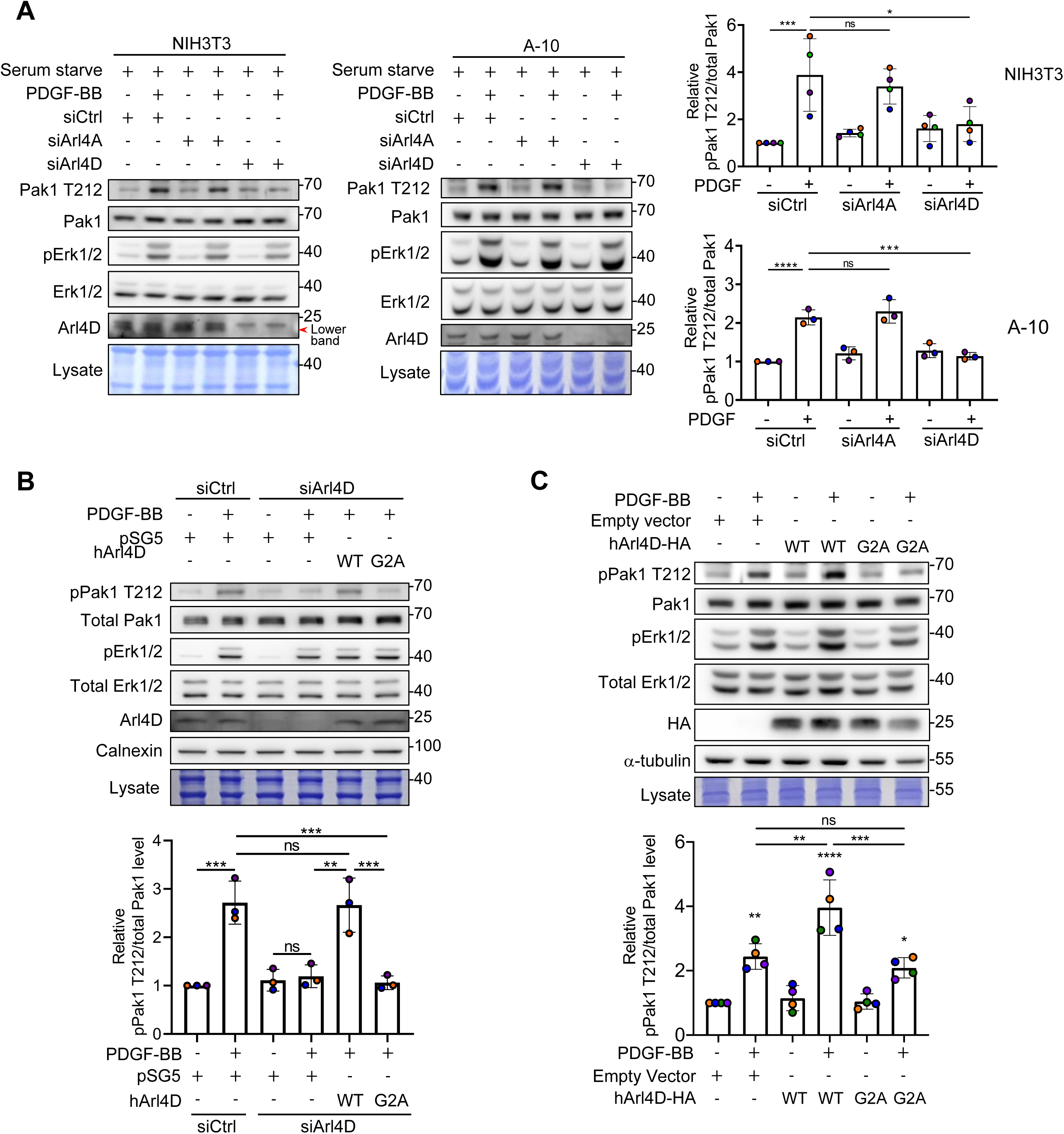
Erk1/2 dependent phosphorylation of Pak1 is rely on membrane-bound Arl4D. (A) NIH3T3 and A10 cells, after knocked down with siCtrl, mouse siArl4A, or mouse siArl4D RNA, were serum starved and treated with PDGF-BB (20 ng/mL) for 10 min. Cell lysates immediately harvested were subjected to SDS-PAGE and Western blotting for indicated proteins. The red arrow indicates the position of Arl4D. The pPak1 T212 signals were divided with total Pak1 signals and normalized with the value of the first control group. The quantified results are shown in the dot plots with error bars indicating the mean ± SD (NIH/3T3, n=4; A-10, n=3). *P<0.05; ***P<0.001; ****P<0.0001 (one-way ANOVA with Tukey’s post hoc multiple comparison test). (B) NIH3T3 cells knocked down with siCtrl or mouse siArl4D RNA and transfected with empty vector pSG5 or human Arl4D-WT or Arl4D-G2A were serum starved and treated with PDGF-BB (20 ng/mL) for 10 min. Cell lysates immediately harvested were subjected to Western blotting for indicated proteins. The pPak1 T212 signals were divided with the total Pak1 signals and normalized with the value of the first control group. The quantified results are shown in the dot plots with the error bars indicating the mean ± SD (n=3). **P<0.01; ***P<0.001 (one-way ANOVA with Tukey’s post hoc multiple comparison test). (C) NIH3T3 cells transfected with empty vector pSG5, Arl4D-WT-HA or Arl4D-G2A-HA were serum starved and treated with PDGF-BB at 20 ng/mL for 10 min. Immediately harvested cell lysates were subjected to Western blotting for indicated proteins. The pPak1 T212 signals were divided with total Pak1 signals and normalized with the value of the first control group. The quantified results are shown in the dot plots with the error bars indicating the mean ± SD (n=4). **P<0.01; ***P<0.001; ****P< 0.0001 (one-way ANOVA with Tukey’s post hoc multiple comparison test).

### Arl4D recruits Erk1/2 to the plasma membrane upon PDGF stimulation

After extracellular stimuli trigger Erk1/2 activation, regulator(s) are required to facilitate the transient translocation of activated Erk1/2 to its target substrates (Ebisuya *et al*, 2005; Kolch, 2005). A notable finding above was that Arl4D did not affect Erk1/2 activation (Fig. 2A, B). Thus, we sought to gain a better temporal and spatial understanding of Arl4D-Erk1 interaction after PDGF treatment. The IF and co-IP data showed that plasma membrane colocalization and their association increased significantly after PDGF stimulation (Fig. EV2 A, B). We noted that the GTP-locked membrane-bound form of Arl4D (Q80L) could not interact with Erk1/2 in the absence of PGDF stimulation (Fig. EV2B), suggesting that Arl4D binds preferentially to the activated forms of Erk1/2. Further characterizing their interaction, an increase in the co-translocation of Erk1 and Arl4D at the plasma membrane by 10 min after PDGF treatment is shown (Fig. 3A), which is consistent with the previous finding that pPak1 T212 reached peak at 10 min upon PDGF stimulation (Sundberg-Smith *et al*., 2005b). Consistent with this finding, Arl4D interacts transiently with activated Erk1/2 also peaked at about that time (Fig. 3A). Moreover, the association of Pak1 with the co-precipitated Arl4D and Erk1/2 also peaked at 10 minutes (Fig. 3B). To further substantiate Arl4D could be the regulator that directs Erk1/2 to phosphorylate membrane substrates upon extracellular signaling, we found that Arl4D overexpression caused a significant shift of endogenous Erk1/2 from the cytosolic fraction to the membrane fraction upon PDGF treatment (Fig. 3C). Moreover, PDGF-triggered Erk1/2-Arl4D association is much stronger in the membrane fraction than in the cytosolic fraction (Fig. 3D). Thus, the collective results suggested that Arl4D could be acting to bring Erk1/2 and Pak1 together upon PDGF stimulation.

**Fig. 3.**
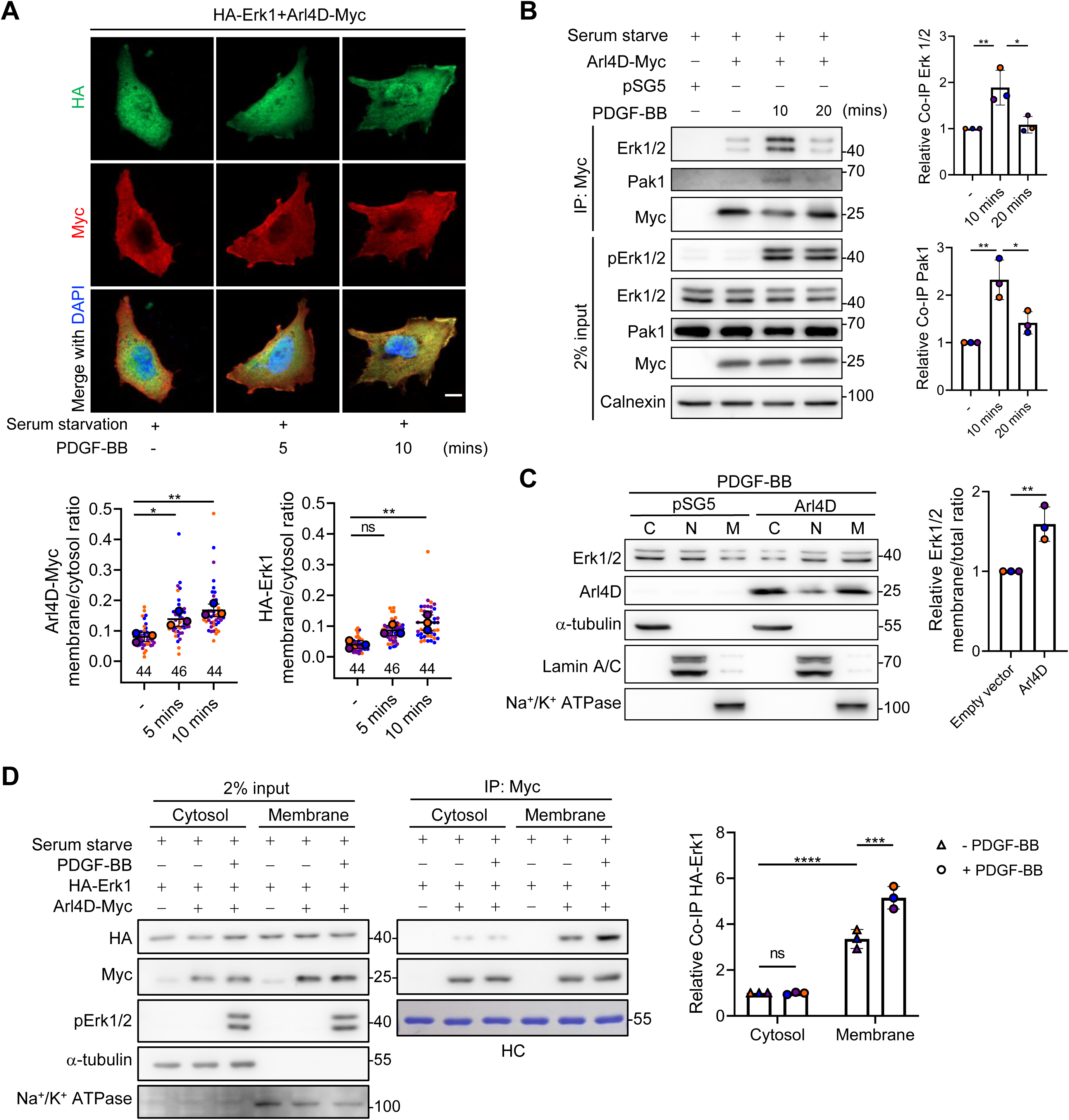
Arl4D complexes and recruits Erk1/2 to plasma membrane under PDGF signaling. (A) NIH3T3 cells transfected with indicated proteins were serum starved, treated with PDGF-BB at 20 ng/mL for 0, 5, 10 min and stained with anti-Myc (red), and anti-HA (green) antibodies and DAPI (blue; stains the nuclei). Scale bar, 10 mm. The ratios between plasma membrane and cytosol of HA-Erk1 in each group were quantified as described in the Materials and Methods, and the results were shown in the dot plots with error bars indicating the mean ± SD (n=3, the cells analyzed in each biological replicate were marked in the same color, and the total number of cells is indicated in the plot). **P<0.01 (one-way ANOVA with Tukey’s post hoc multiple comparison test). (B) NIH3T3 cells transfected with empty vector pSG5 or Arl4D-Myc were serum starved and treated with PDGF-BB at 20 ng/mL for the indicated minutes and subjected to DSP crosslinker-treated co-IP. The Co-IP signals of endogenous Pak1 and Erk1/2 were normalized with Arl4D-Myc proteins after subtracting the background signal from pSG5 control group and the quantified results are shown in the dot plots with error bars indicating the mean ± SD (n=3). *P<0.05; **P<0.01 (one-way ANOVA with Tukey’s post hoc multiple comparison test). (C) NIH3T3 cells transfected with empty vector pSG5 or Arl4D were serum starved, treated with PDGF-BB (20 ng/mL) for 10 min and subjected to cellular fractionation for Western blotting of the indicated proteins. α-tubulin, a cytosolic fraction protein; Lamin A/C, a nuclear fraction protein; Na+/K+ ATPase, a membrane fraction protein. The percentages of endogenous Erk1/2 membrane fraction divided by the total fraction were quantified and normalized with the pSG5 control group. The dot plots with error bars show the mean ± SD (n=3). **P<0.01 (two-tailed unpaired Student’s t-test). (D) NIH/3T3 cells transfected with empty vector pSG5 or Arl4D-Myc and HA-Erk1 were serum starved, treated with PDGF-BB for 10 min and subjected to DSP crosslinker-treated fractionation Co-IP. α-Tubulin, cytosolic fraction protein; Na+/K+ ATPase, membrane fraction protein. The Co-IP signals of HA-Erk1 in were normalized with Arl4D-Myc proteins after subtracting the background signal from pSG5 control group and the quantified results are shown in the dot plots with error bars indicating the mean ± SD (n=3). ***P<0.001, ****P < 0.0001 (one-way ANOVA with Tukey’s post hoc multiple comparison test). HC, heavy chain of the antibody.

### Identification of Erk1/2 mutant defective in binding to Arl4D

To gather more definitive support for this role of Arl4D, we next sought to generate Erk1/2 mutants that cannot interact with Arl4D. We initially focused on generating Erk1 truncations that targeted the disordered region (ΔD), the entire N-terminus (ΔD + ΔN), or the C-terminus (ΔC). Incubating these constructs as purified proteins with recombinant Arl4D in pulldown experiments, we found that the entire N-terminus of Erk1 (ΔD + ΔN) is critical for its direct interaction with Arl4D (Fig. EV3A). Further dissecting out this region, we found that only aa 27-41 (designated as ΔN) of Erk1 is required for its interaction with Arl4D (Fig. EV3B).

The region in Erk2 that corresponds to aa 27-41 in Erk1 spans residues 11-24 (Fig. EV4A). Consistent with the finding that ΔN serves as a critical binding site for Arl4D, both Erk1and Erk2 that lacks this region (ΔN) showed impaired co-localization to the plasma membrane (Fig. EV4B). This suggests that Arl4D-Erk1/2 interaction is essential for Erk1/2 plasma membrane targeting. Furthermore, we confirmed that both Erk1/2 ΔN mutants were still responsive to PDGF stimulation, as assessed by their phosphorylation (p-ERK1/2) status upon stimulation (Fig. EV4C).

### PDGF-induced targeting of Pak1 by Erk1/2 requires Arl4D

With the Erk1/2 mutants (ΔN) in hand, we then examined whether the ability of Erk1/2 to associate with Pak1 requires Arl4D. We initially immunoprecipitated for Erk1/2 in cell lysates and confirmed that the WT forms of Erk1/2 associated with both Arl4D and Pak1 (Figs. 4A and EV4D). In contrast, the mutant (ΔN) forms of Erk1/2 not only did not associate with Arl4D, but also lost association with Pak1 (Figs. 4A and EV4D). We next immunoprecipitated for Pak1 and found that PGDG treatment stimulated the association of Pak1 with activated Erk1/2, while siRNA against Arl4D prevented this association (Fig. 4B). We also performed IF and found that siRNA against Arl4D reduced the PDGF-induced Erk1 plasma membrane targeting and colocalization of Erk1 with Pak1 (Fig. 4C). Thus, these results provided more definitive support that Arl4D is critical for Erk1/2 to associate with Pak1.

**Fig. 4.**
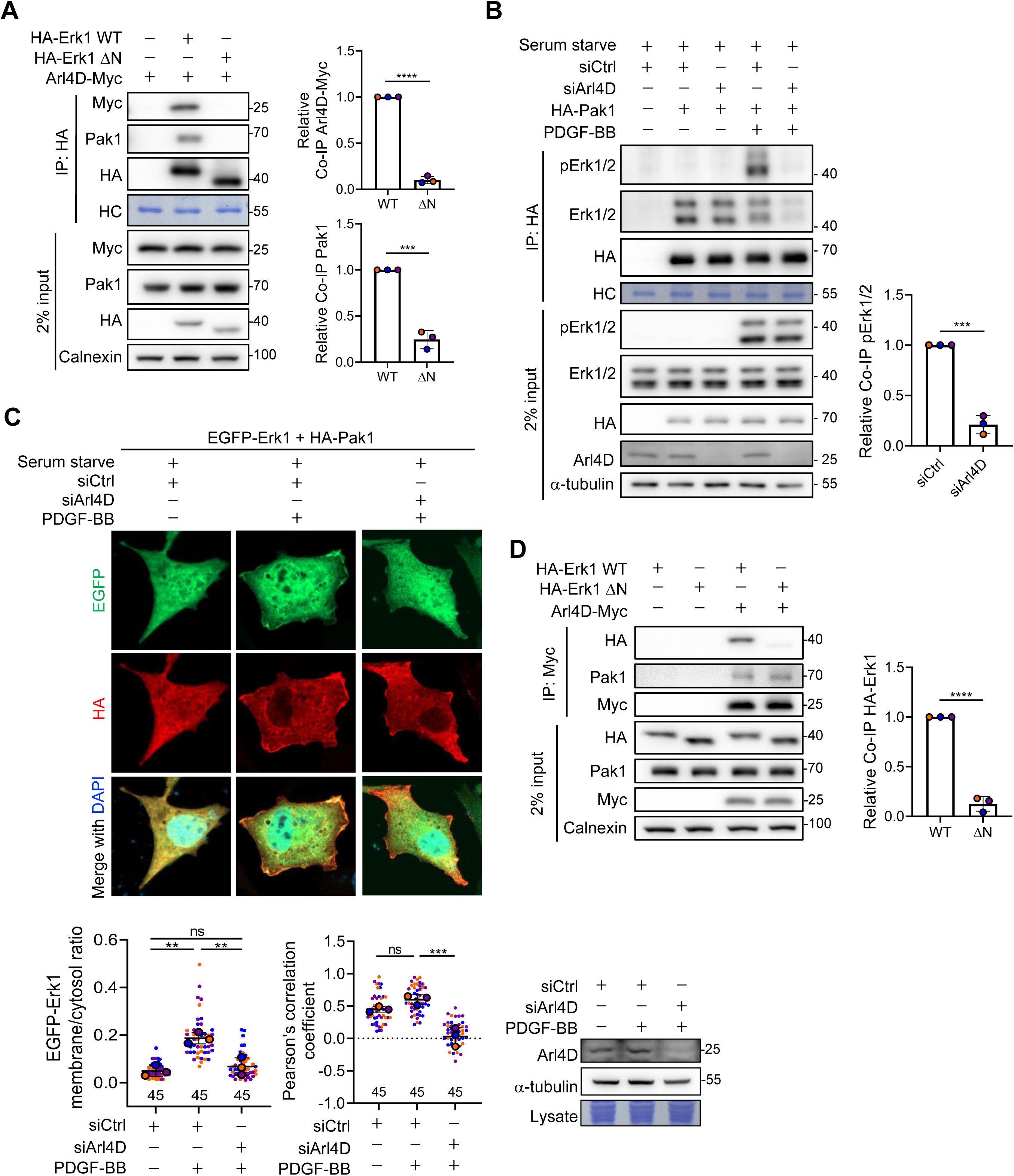
Erk1/2 requires Arl4D to target Pak1 under PDGF signaling. (A) NIH3T3 cells transfected with empty vector pcDNA3.0-HA, Arl4D-Myc, HA-Erk1 WT or HA-Erk1 ΔN were subjected to Co-IP. The Co-IP signals of endogenous Pak1 and Arl4D-Myc were normalized with HA-Erk1 proteins after subtracting the background signal of the empty vector control group. HC, heavy chain of the antibody. (B) NIH3T3 cells knocked down with siCtrl or mouse siArl4D RNA and transfected with empty vector pcDNA3.0-HA or HA-Pak1 were serum starved, treated with PDGF-BB (20 ng/mL) for 10 min, and subjected to DSP-crosslinker-treated Co-IP. The Co-IP signals of endogenous pErk1/2 were normalized with HA-Pak1 proteins after the background signal of pcDNA3.0-HA group was subtracted. HC, heavy chain of the antibody. (C) NIH3T3 cells transfected with the indicated proteins were stained with anti-HA (red) antibodies and DAPI (blue; stains the nuclei). Scale bar, 10 mm. Pearson correlation coefficients of EGFP-Erk1 and HA-Pak1 signals were calculated using ZEN imaging software, respectively, and the results were shown in the dot plots with error bars indicating the mean ± SD (n=3, cells analyzed in each biological replicate were marked in the same color, and the total number of cells is indicated in the graph). The ratios between plasma membrane and cytosol of GFP-Erk1 in each group were quantified as described in the Materials and Methods, and the results were shown in the dot plots with error bars indicating the mean ± SD (n=3, the cell samples are the same as described). *P<0.05; **P<0.01 (one-way ANOVA with Tukey’s post hoc multiple comparison test). Western blotting demonstrated the knockdown efficiency of Arl4D. (D) NIH3T3 cells transfected with empty vector pSG5, Arl4D-Myc, HA-Erk1 WT or HA-Erk1 ΔN were subjected to co-IP. The co-IP signals of HA-Erk1 were normalized with Arl4D-Myc proteins after subtracting the background signal of the control group with empty vector. (A-B & D) The quantified results are shown in the dot plots, with the error bars indicating the mean ± SD (n=3). ***P<0.001, ****P < 0.0001 (two-tailed unpaired Student’s t-test).

We next examined whether Erk1/2 or Pak1 may influence the ability of each other to interact with Arl4D. Upon immunoprecipitating for Arl4D, we first confirmed that Arl4D associated with both Erk1/2 WT and Pak1 (Figs. 4D and EV4E). However, whereas Arl4D did not associate with the Erk1/2 mutants (ΔN), as expected, it retained association with Pak1 (Figs. 4D and EV4E). Moreover, performing siRNA against Pak1, we found that this treatment had no appreciable effect on the Arl4D-Erk1/2 interaction (Fig. EV5A), as well as the recruitment of Erk1/2 to the plasma membrane by Arl4D (Fig. EV5B). Thus, neither Erk1/2 nor Pak1 regulates the ability of the other to associate with Arl4D.

### The interplay among Arl4D, Erk1/2 and Pak1 enhances cell migration

We next considered that the biological significance of Erk1/2 phosphorylating Pak1 at its T212 residue with respect to cell migration remains to be better established. Thus, we first generated phosphorylation mutants of the T212 residue, with mutation to alanine (T212A) to abrogate phosphorylation and mutation to aspartate (T212D) to mimic phosphorylation. Upon siRNA against Pak1 followed by rescues using these phosphorylation mutants, we found that the expression of the T212A mutant reduced the migration of NIH3T3 cells while the expression of the T212D mutant enhanced migration (Fig. 5A, D). Consistent with these findings, cell migration that was enhanced by Pak1 overexpression was further promoted by PDGF treatment, and in this setting, we observed enhanced Erk1/2 activation and Pak1 phosphorylation at T212, which were prevented by U0126 treatment (Fig. EV7A). Moreover, whereas siRNA against Erk1/2 reduced cell migration, the expression of the Pak1 T212D mutant bypassed this inhibition (Fig. EV7B), and thus further confirming that Pak1 acts downstream of Erk1/2 for cell migration.

**Fig. 5.**
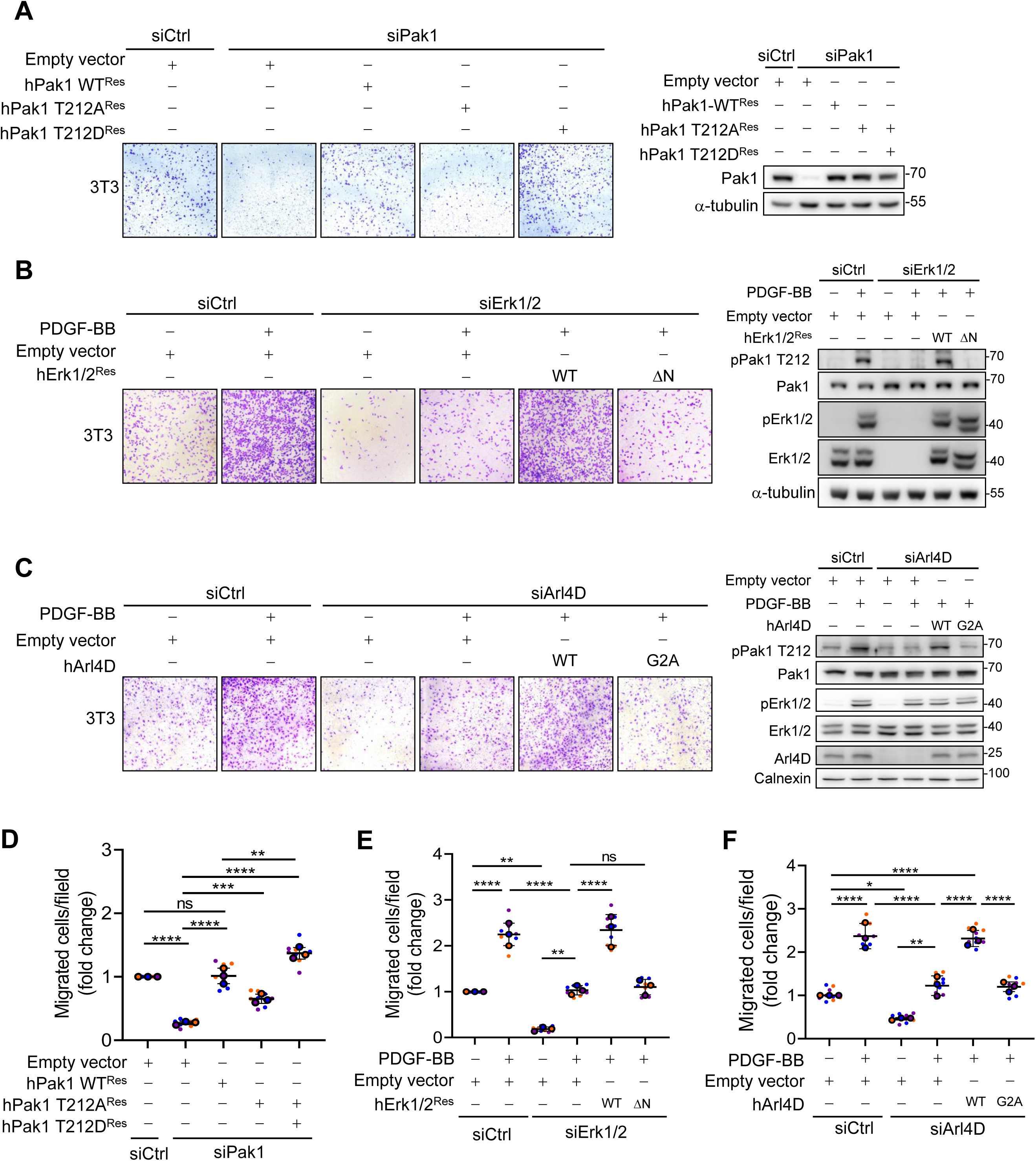
Arl4D plays a pivotal role in PDGF-Erk1/2-Pak1 mediated cell migration. (A) NIH3T3 were knocked down with siCtrl or mouse siPak1 RNA and transfected with empty vector pcDNA3.0, hPak1-WT^Res^, hPak1-T212A^Res^ or hPak1-T212D^Res^. The cells were subjected to the normal conditions cell migration as material and methods described. (B) NIH3T3 were knocked down with siCtrl or mouse siErk1/2 RNA and transfected with empty vector pcDNA3.0 or hErk1/2^Res^WT, hErk1/2^Res^ΔN. (C) NIH3T3 were knocked down with siCtrl or mouse siArl4D RNA and transfected with empty vector pSG5 or human Arl4D-WT or Arl4D-G2A. (B and C) The cells were subjected to the PDGF conditions cell migration as material and methods described. (A-C) Western blotting detected the indicated proteins. (D-F) Cells migrated under the transwells were stained with crystal violet and photographed for 3 fields per biological replicate and calculated in dot plot with mean ± SD (n=3). *P<0.05; **P<0.01; ***P<0.001; ****P< 0.0001 (one-way ANOVA with Tukey’s post hoc multiple comparison test).

Having established the critical role that the T212 phosphorylation of Pak1 plays in cell migration, we next examined how the interplay among Arl4D, Erk1/2, and Pak1 that we have elucidated affects cell migration. When endogenous Erk1/2 was replaced by the Erk1/2 mutant that could not bind Arl4D (ΔN), we found that PDGF treatment could no longer enhance cell migration (Fig. 5B, E). Further confirming the critical role of Arl4D, we depleted its level by siRNA treatment followed by rescue using the myristoylation-defective mutant of Arl4D (G2A), and found that PDGF stimulation could no longer enhance cell migration (Fig. 5C, F). It is further notable that the phosphorylation of T212 in Pak1 was inhibited in this setting, while Erk1/2 activation was unaffected (Fig. 5C). These additional observations provided further support that the scaffolding role of Arl4D, which brings Erk1/2 and Pak1 together, is critical for cell migration.

## Discussion

Whereas Arl4D, Erk1/2, and Pak1 have all been found to act in cell migration, how the interplay among them affects cell migration has been unknown. Our study addresses this outstanding question by elucidating that Arl4D acts as a scaffolding protein that enables activated Erk1/2 to target Pak1 for phosphorylation at its T212 residue, resulting in cell migration being promoted (Fig. 6). This elucidation also addresses a more general question. Scaffolding proteins are known to be needed for activated Erk1/2 that translocate from the plasma membrane to other intracellular site in targeting substrates. However, it has been less clear whether scaffolding is also needed when activated Erk1/2 remain at the plasma membrane in targeting their substrates. Our finding that Arl4D acts as scaffolding protein so that activated Erk1/2 can target Pak1 at the plasma membrane suggests that scaffolding is a conserved requirement for substrate targeting by Erk1/2.

**Fig. 6.**
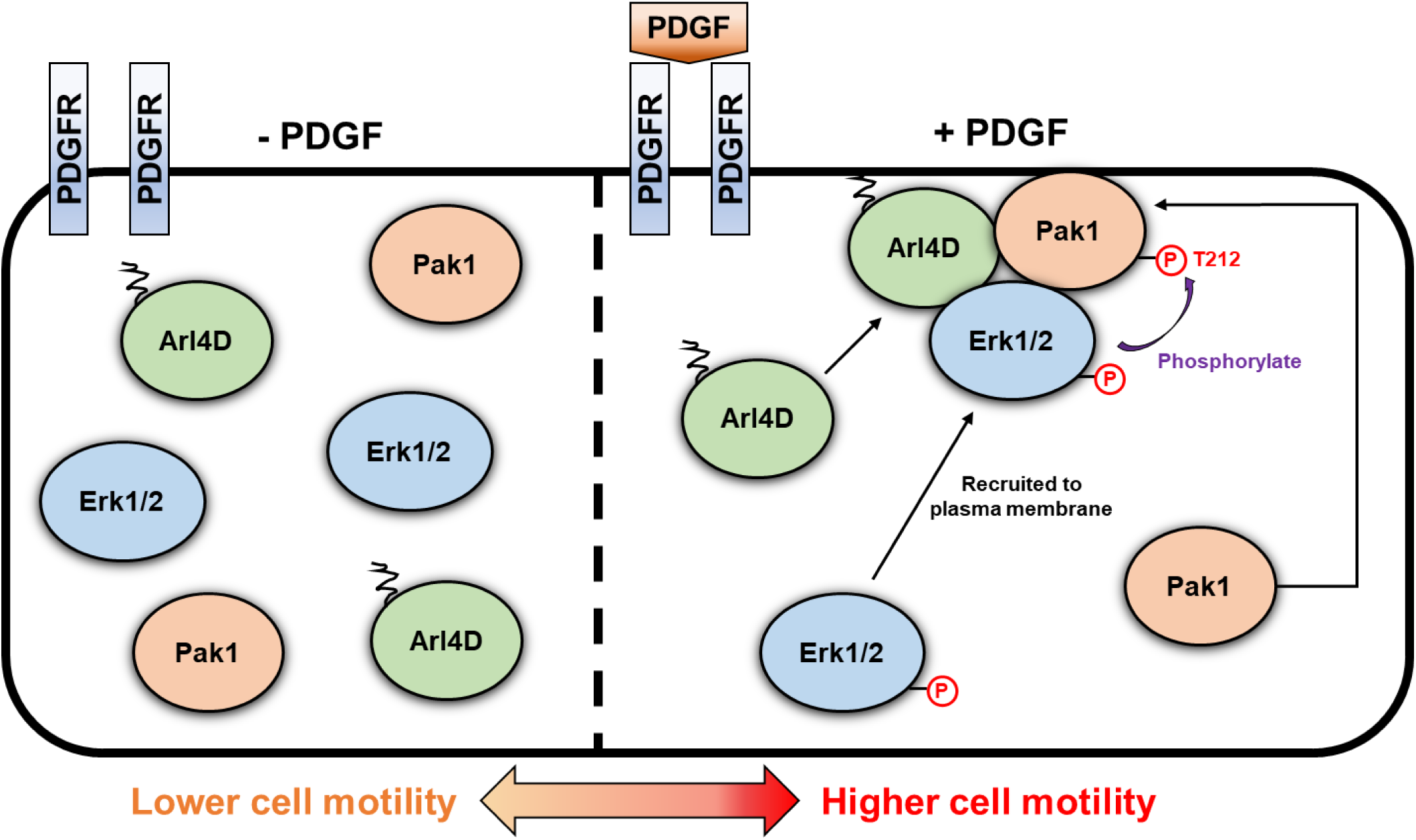
The mechanistic model of PDGF-driven cell migration through Arl4D-Erk1/2-Pak1 interaction. Under PDGF treatment, Arl4D plays the important role in interacting and recruiting activated Erk1/2 as well as Pak1 to the plasma membrane. Following, Arl4D serves as the interbridge between activated Erk1/2 and Pak1 to make Pak1 T212 motif be phosphorylated successfully, which promote the cell motility.

The overall complexity of cell migration involves various signaling mechanisms, including PDGF-induced Erk1/2 activation, the interplay with GPCRs (guanine nucleotide-binding protein-coupled receptors), ECM (extracellular matrix), and integrins (Abedi & Zachary, 1995; Hobson *et al*, 2001; Veevers-Lowe *et al*, 2011). Our previous study has shown that fibronectin, an ECM component, stabilizes Arl4A/D protein levels at the plasma membrane and triggers Pak1 activation (Chen *et al*, 2020b; Lin *et al*, 2022b). Nevertheless, fibronectin has a very mild effect on Pak1 T212 phosphorylation (Sundberg-Smith *et al*., 2005b). This reminds us that functions of a protein in promoting cell migration could be underestimated if explained by a single signaling cue.

Our results also address another puzzle. We had found previously that Arl4D and Pak1 recruit each other to the plasma membrane, which is mediated by their interaction. However, a further finding was that this recruitment results in the phosphorylation in Pak1, but the identity of a predicted kinase had remained elusive (Chen *et al*., 2020a). Our results now reveal Erk1/2 as the responsible kinases. We further note that Erk1/2 and Pak1 are involved in complex feedback loops that regulate the activities of each other. Phosphorylation of Pak1 at T212 has been found to decrease Erk1/2 activation (Pullikuth *et al*, 2005; Sundberg-Smith *et al*., 2005a). Moreover, Pak1 has been found to phosphorylate MEK1, which also leads to decreased Erk1/2 activation (Beeser *et al*, 2005; Park *et al*, 2007). As such, it has been unclear what specific circumstance might enhance the phosphorylation of Pak1 at T212 and lead to enhanced cell migration. We define one such circumstance through the interplay that we have elucidated among Arl4D, Erk1/2, and Pak1.

## Materials and methods

### Cell culture

The cell lines C33A (HTB-31), A549 (CCL-185), Hela (CCL-2), Cos7 (CRL-1651), NIH3T3 (CRL-1658), A-10 (CRL-1476), TM3 (CRL-1714) and TM4 (CRL-1715) were purchased from the American Type Culture Collection (ATCC, Manassas, VA, USA). Cell lines TM3 and TM4 were maintained in F12 medium and Dulbecco’s modified Eagle’s medium (DMEM) (1:1) (Gibco™ 11330032) with 2.5 mM L-glutamine, 0.5 mM sodium pyruvate, 1.2 g/L sodium bicarbonate and 15 mM HEPES, 5% horse serum, and 2.5% fetal bovine serum (FBS) (Biological Industry). Other cell lines was maintained in high glucose DMEM (Cytiva, SH30003.03) with 1mM sodium pyruvate, 1.5 g/L sodium bicarbonate, 10% FBS (35-010-CV, Corning) and pH 7.4. For A-10, 4 mM L-glutamine was supplied. All cell lines were maintained in a humidified incubator with 5% CO_2_ at 37°C.

### Antibodies

For Western blotting, the following primary antibodies were used at the indicated dilutions: anti-α-tubulin (1:10000, Sigma), anti-His (1:5000, #631212, Takara), anti-Erk1/2 (1:2000, #4695S, Cell Signaling), anti-pErk1/2 (1:2000, #4370S, Cell Signaling), anti-Pak1 (1:2000, #2602S, Cell Signaling), anti-pPak1 T212 (1:1000, #PA5-37677, Invitrogen), anti-Myc (1:3000, #2278S, Cell Signaling), anti-HA (1:3000, #3724S, Cell Signaling), anti-Na^+^ /K^+^ ATPase (1:1000, #3010S, Cell Signaling), anti-Lamin A/C (1:2000, #4777S, Cell Signaling), anti-Arl4D (1:1000, boosted from our lab), anti-GFP (1:1000, boosted from our lab), anti-Calnexin (1:1000, boosted from our lab) and anti-ϕ3-actin (1:2000, #GTX629630, GeneTex). The secondary antibodies were goat horseradish peroxidase (HRP)-conjugated anti-rabbit/mouse IgG (1:5000, NA934V/NA931V, GE Healthcare). For the immunofluorescence analyses, dilutions of primary antibodies anti-HA (1:200, #901515, Biolegend), anti-Myc (1:200, #2278S, Cell Signaling), and anti-Arl4D (1:400, boosted from our lab) were used. The secondary antibodies were Alexa Fluor 488/594 goat anti-rabbit and mouse IgG secondary antibodies (1:1000, A-31556/A-11034/A-11032, Invitrogen). The DAPI (4’,6-diamidino-2phenylindole) solution was purchased from Millipore (1:5000, S7113).

### Plasmid DNAs and siRNAs

In the mammalian expression system, HA- or Myc-tagged Arl4A, Arl4C, Arl4D and non-tagged Arl4D and Arl4D-G2A mutants, were cloned into the pSG5 vector (Stratagene) as previously described (Chen *et al*., 2020a). Erk1, Erk2 and their truncations (NC_000022.11) derived from C33A cell cDNA library were cloned into pcDNA3.1 (Invitrogen), pcDNA3.0-HA (Invitrogen) or pEGFP-C3 vectors (Clontech). Human Pak1 in pCMV6M-HA were kindly provided by professor Jau-Song Yu. Pak1 WT, Pak1 T212A and Pak1 T212D were subcloned into the pcDNA3.0 vector (Invitrogen). For the bacterial expression system, Erk1, Erk2 and Arl4D were subcloned into pGEX-4T-1 (GE Healthcare) vectors to obtain the N-terminal GST-fusion proteins. Arl4A, Arl4C and Arl4D, as well as the GTP-bound and GTP-deficient mimetic mutants of Arl4Dwere subcloned into the pET15b (Novagen) vectors to obtain the N-terminal His-tagged proteins. Erk1 and its truncations were subcloned into pET30a vector (Novagen, Madison, WI).

Pak1 WT^Res^, Erk1^Res^ and Erk2^Res^ siRNA resistant constructs were generated by site-directed mutagenesis with the following primers: (F: 5’-GAGCACACAAAATCTGTCTACACGCGGTCCGTTATTGAACCACTTCC-3’, R: 5’-GGAAGTGGTTCAATAACGGACCGCGTGTAGACAGATTTTGTGTGCTC-3’); (F: 5’-CTCCCCATCCCAGGAGGATCTGAACTGTATTATCAACATGAAGGCCC-3’, R: 5’-GGGCCTTCATGTTGATAATACAGTTCAGATCCTCCTGGGATGGGGAG-3’); (F: 5’-GACATTCAACCCACATAAAAGAATCGAGGTCGAACAGGCTCTGGCC, R: 5’-GCCAGAGCCTGTTCGACCTCGATTCTTTTATGTGGGTTGAATGTC-3’). The Pak1 T212A^Res^ and T212D^Res^ site-directed mutagenesis used Pak1 WT^Res^ as the template and PCR with the following primers respectively: (F: 5’-GAACCACTTCCTGTCGCTCCAACTCGGGACGTG-3’, R: 5’-CACGTCCCGAGTTGGAGCGACAGGAAGTGGTTC-3’); (F: 5’-GAACCACTTCCTGTCGATCCAACTCGGGACGTG-3’, R: CACGTCCCGAGTTGGATCGACAGGAAGTGGTTC-3’).

All mouse siRNAs were purchased from Dharmacon and the sequences used in this paper are as follows: siArl4A: AAAGGAAGGACUCGAGAAA; siArl4D: GAAUGGAGUUACACCGGAU; siPak1: GUAUAUACACGAUCUGUGA; siErk1: GGACCUUAAUUGCAUCAUU; siErk2: ACAAGAGGAUUGAAGUUGA; siCtrl: UGGUUUACAUGUCGACGACUAAUU.

### Transient expression and knockdown, total protein extraction and Western blotting

Cells were transiently transfected with the indicated plasmids using Lipofectamine 2000 or siRNAs using RNAiMax transfection reagent (Invitrogen) in Opti-MEM (Gibco), following the manufacturer’s instructions.

After transfection and PDGF or drug treatment, cells were scraped and lysed with RIPA buffer (50 mM Tris-HCl, pH 7.5; 0.1% sodium dodecyl sulfate; 0.5% sodium bicarbonate; 1% NP-40; 150 mM NaCl and 1 mM EDTA) containing protease inhibitor (PI) cocktail and PhosSTOP™ (04906837001, Roche). The cell lysates were centrifuged at 15000 g for 10 minutes to remove insoluble debris. The supernatant was mixed with 5-fold sample buffer (250 mM Tris-HCl, pH 6.8; 10% sodium dodecyl sulfate; 50% glycerol; 0.1% bromophenol blue and 5% 2-mercaptoethanol) and boiled at 95°C for 10 minutes before subjecting to 10% SDS-PAGE gels. Western blotting analysis was performed according to previously established protocols (Chen et al., 2020). 0.45-μm PVDF membranes (Immobilon P, Millipore) were used for protein transfer and Tident femto Western HRP Substrate (GTX14698) was applied for chemiluminescence system (ImageQuant™ LAS 4000). α-tubulin, ϕ3-actin or calnexin were used as an internal control for protein loading. GST-tagged and His-tagged proteins were visualized by Coomassie blue staining.

### PDGF and drug treatment

After transfection of the indicated plasmids or siRNA, the cells were washed twice with PBS and then supplied with serum-free culture medium (serum starvation) for 16-24 hr. Serum-free medium containing 20 ng/ml platelet-derived growth factor-BB (PDGF-BB) (Z02892, GenScript) was used to trigger Erk1/2 activation. To block Erk1/2 activation, 10 uM MEK inhibitor U0126 (V112A, Promega) was co-treated with PDGF-BB.

### RNA extraction and RT-PCR

To determine the mouse Arl4A mRNA expression level in NIH3T3 and A-10 cells, total RNA was extracted using the GENEzol Pure Kit (GZXD200, Geneaid) and subjected to an in-column DNase I digestion procedure. 1 μg of total RNA was reverse transcribed (RT) using the MMLV Reverse Transcription Kit (PT-RT-KIT, PROTECH). Subsequently, 1 μL of the RT product was used to perform polymerase chain reactions (PCRs) using VeriFi^TM^ Mix (PB10.43-02, PCRBIOSYSTEMS), with mouse Arl4A specific primers for 25 cycles and with GAPDH primers for 20 cycles. The PCR products were loaded in a 1.5% agarose EtBr gel for UV detection. The following PCR primers were used: Arl4A (F, 5’-AGAGCGGCGCGGAGGTCTCGGTTGAG-3’, and R, 5’-GGTACGGTATTTACAAATTCGTTGAAC-3’); and GAPDH (F, 5’-TATTGGGCGCCTGGTCACCAG-3’, and R, 5’-GAGATGATGACCCTTTTGGCTCC-3’).

### Protein purification and *in vitro* binding assay

*E. coli* expression plasmids for production of glutathione S-transferase (GST), and GST fusion proteins, Erk1, Erk2 and their truncations as well as His-tagged Erk1, its truncations and Arl4s proteins were transformed into competent BL21 (DE3) cells. Colonies expressing the recombinant proteins were cultured at 37 °C in LB medium until they reached log phase (OD600=0.6-0.8). Isopropyl-β-D-1-thiogalactopyranoside (IPTG) and the specific induction conditions were then applied as follows: Erk1, Erk2 and their truncations, and Arl4D were induced with 0.5 mM IPTG at 25°C for 3 hr, Arl4A and Arl4C were induced with 0.5 mM IPTG at 37°C for 3 hr.

For purification of GST fusion proteins, cell pellets were lysed in PBS with a protease inhibitor (PI) cocktail and 0.5% Triton X-100. For purification of His-tagged proteins, the cell pellets were lysed in a lysis buffer (20 mM Tris–HCl, pH 7.9; 500 mM NaCl; 5 mM imidazole; 10 % glycerol; 0.1 % Triton X-100; PI cocktail; and 100 μg/mL lysozyme). Suspensions were incubated on a 3D shaker for 30 minutes at 4 °C and protiens were purified as previously described (Lin *et al*., 2022b). Alternatively, the His-tagged proteins were eluted with elution buffer (20 mM Tris-HCl, pH 7.9; 500 mM NaCl; and 500 mM imidazole).

Quantification of GST fusion proteins or His-tagged proteins was performed by SDS-PAGE with BSA (11930.03, SERVA) as standards and followed by Coomassie blue staining. The protein binding processes were as mentioned before (Lin *et al*., 2022b).

### Co-immunoprecipitation (Co-IP)

If not alternatively specified, cells for Co-IP were lysed on ice with 1 ml of lysis buffer (50 mM Tris-HCl, pH 7.4; 150 mM NaCl; 1 % NP-40; 1 mM EDTA; 5 % glycerol; 0.1 % Tween-20; and a protease inhibitor (PI) cocktail) per 10-cm dish. Lysates were then rotated at 4°C for 30 minutes followed by centrifugation at 15000 g for 10 minutes to remove insoluble debris. Pierce™ Anti-HA Magnetic Beads (88836, Thermo Scientific), Myc-Trap Magnetic Agarose (ytma-20, ChromoTek) or GFP-Trap Magnetic Agarose (gtma-20, ChromoTek) were pre-washed three times with lysis buffer and then incubated with the supernatants on an end-to-end rotator at 4°C for 2 hours. After, the beads were washed three times with lysis buffer over a Magna GrIP rack (Millipore) and resuspended in 30 μl of 1X sample buffer. The eluted protein complexes were then boiled at 95 °C for 10 minutes prior to Western blotting analysis.

For Co-IP after PDGF-BB treatment, cells were treated with 100 mM dithiobis-[succinimidyl propionate] (DSP; 22585, Thermo Scientific) for cross-linking for 2 hrs, according to the well-described protocol (Zlatic et al, 2010). After quenching and washing, the supernatant was exposed to prewashed Myc-Trap magnetagarose (ytma-20, ChromoTek) in an end-over-end rotator at 4°C for 2 hours. The beads were then washed three times with lysis buffer followed by resuspension of 30 μl of 1X sample buffer containing 0.2 M DTT. The sample was then incubated at 37 °C for 1 hour and then boiled at 95 °C for 10 min prior to Western blotting analysis.

For Co-IP cytosol and membrane fractionation, cells were DSP cross-linked as mentioned above. Subsequent membrane and cytosol fractionation and Co-IP protocols are following as established (Lin *et al*., 2022b).

### CNM fractionation

Following serum starvation and PDGF-BB treatment, NIH3T3 cells (1×10^7^) were washed twice with PBS. Subsequently, 500 μL of ice-cold buffer C (K3012010, Biochain Institute) was added to scrape and resuspend the cell pellet. The suspension was rotated at 4°C for 20 minutes. The cells were homogenized by passing them 25 times through a 30-gauge needle, resulting in the release of 90–95% of the cell nuclei, as determined by microscopy. The following protocol applying Buffer W, N, and M (K3012010, Biochain Institute) is used according to the manufacturer. All fractionated proteins were mixed with 5X sample buffer and denatured at 60°C for 15 minutes before Western Blotting analysis.

### Immunofluorescence (IF) staining

After transfection and treating with or without PDGF, the coverslips were washed twice with PBS and then fixed in 4% paraformaldehyde in PBS for 15 minutes at room temperature. After following the protocol as described (Lin *et al*., 2022b), fluorescence images were captured with an AxioImager.Z2 microscope with ApoTome.2 (Carl Zeiss, Inc.) and analysed with FIJI software (ImageJ2). ZEN software is used to determine the colocalization coefficient of two proteins of interest. To evaluate the plasma membrane targeting of interested protiens, the fluorescence intensity of the plasma membrane to the cytosol (PM:C) the ratio was calculated using the following formula: PM/C ratio = (intensity of cell body area-cytosol area)/intensity of cytosol area. The quantification methods have been described previously (Barry et al., 2015; Chen et al., 2020).

### Transwell migration assay

Transwell devices (3464, Corning) were coated with FN (10 μg/mL, 356008, Corning) in PBS in 24-well culture plates (142475, Thermo Scientific) for 2 hrs. After knockdown or/and transient transfection, cells which had been through serum starvation for 16-18 hours were counted. 5×10^4^ NIH3T3 cells were resuspended in 200 μl serum-free culture medium and seeded into the Transwell device, with bottom supplied with medium containing 10% FBS. For cell migration under PDGF conditions, the serum-free culture medium containing 20 ng/ml PDGF and/or 10 μM MEK inhibitor U0126 were added. The quiescent cells were scraped with RIPA buffer for Western blotting analysis. The newly seeded cells migrated through the Transwell membrane for 9 hours and were analyzed as previously described (Chen et al., 2020).

### Liposome preparation and floating assay

To prepare liposomes, lipids (Avanti Polar Lipids, Inc) in chloroform were mixed in the following ratio: 45 % DOPC, 19 % DOPE, 5 % DOPS, 10 % PI (soybean), 16 % cholesterol (ovine wool), 5 % DSG-NTA(Ni) and 0.2 % NBD-labeled PE. The lipid solvent was evaporated in vacuum machine (IR MICRO-CENVAC) for 1 h and then rehydrated in liposome floating buffer (50 mM HEPES, 120 mM KOAc, 1 mM MgCl_2_, 1 mM DTT) to a final concentration of 2 mg/ml. After freeze and thaw cycle from liquid nitrogen, lipids were filtered through a PC membrane with a pore size of 400 nm using a mini-extruder (Avanti Polar Lipids, Inc.).

200 ul liposomes were incubated with 50 μg purified His-Arl4D Q80L or His-Arl4D T35N in liposome floating buffer containing 200 nM nucleotide (GTP-γ-S for Q80L or GDP for T35N) for 30 min at 30°C. Three layers of sucrose gradients were prepared by adjusting the liposomes to 30% sucrose in liposome floating buffer as the bottom layer, overlaid with 10% sucrose as the middle layer, and free of sucrose at the top. The Arl4D-loaded liposomes were collected from the top layer after centrifugation by SW55Ti rotor at 40 k rpm for 30 mins and then incubated with 50 ug of precleared Hela cell lysate (prepared by Dounce homogenization and sonication in liposome floating buffer and centrifuge at 100,000 g for 2hrs). The mixture was incubated at 4°C for 1 hr and subjected to sucrose gradient centrifuge as described above. The liposome on the top were collected and washed by sucrose gradient centrifuge again, and protein in the top layer were precipitated by methanol and chloroform and dissolved in SDS sample buffer to perform SDS-PAGE.

### Mass spectrometry and Data analysis

Proteins were separated in SDS-PAGE and gel bands were excised followed by in-gel tryptic digestion as described previously (Lin *et al*, 2022a; Liu *et al*, 2015). The peptide mixtures were extracted with acetonitrile to a final concentration of 50% and dried in a SpeedVac. For liquid chromatography-tandem mass spectrometry analysis (LC-MS/MS) analysis, each peptide mixture was resuspended in HPLC buffer A (0.1% formic acid, Sigma) and loaded into a trap column (Zorbax 300 SB-C18, 0.3 × 5 mm, Agilent Technologies), washed with buffer A, and the desalted peptides were then separated on a 10 cm analytical C18 column (inner diameter, 75 mm). The LC setup was coupled to a LTQ-Orbitrap linear ion trap mass spectrometry (Thermo Fisher Scientific). Full-scan MS was performed using the Orbitrap in an MS range of 400–2000 Da and intact peptides were detected at a resolution of 30,000. The data-dependent procedure that alternated between one MS scan followed by 10 MS/MS scans for the 10 most abundant precursor ions in the MS survey scan was applied. For data processing, the resulting mass spectra were searched against the SwissProt_2023 database (Taxonomy: Homo sapiens) using the Mascot search engine (Matrix Science London, U.K.). To identify the proteins, the raw spectrometry data were analyzed using Proteome Discoverer software (version 1.4, Thermo Fisher Scientific). The search parameters were set as follows: trypsin as a digestion enzyme, carbamidomethylation (C) as static modification, oxidation (M), N-acetyl (protein) and Gln->pyro-Glu (N-term Q) as dynamic modification, 10 ppm for MS tolerance, 0.7 Da for MS/MS tolerance, and two for missing cleavage.

### Statistical analysis

All data are presented as mean ± standard deviation (SD) values. P values were calculated using either a two-tailed unpaired Student’s t-test or one-way analysis of variance (ANOVA) followed by Tukey’s post hoc multiple comparison test, performed using Excel or Prism 8 software. Significant differences are denoted as follows: *P ≤ 0.05; **P ≤ 0.01; ***P ≤ 0.001 and ****P ≤ 0.0001. Each independent *in vitro* experiment comprised three biological replicates, and error bars represent independent experiments.

## Author contributions

Ting-Wei Chang, Ming-Chieh Lin, and Fang-Jen S. Lee conceived and designed the research project; Ting-Wei Chang and Ming-Chieh Lin performed the majority of the research and analyzed the data; Chia-Jung Yu supported the experiments and analyzed the data; Ting-Wei Chang and Ming-Chieh Lin drafted the manuscript. All authors edited the manuscript. Fang-Jen S. Lee supervised the project, acquired funding, and performed project administration. All authors read and approved the manuscript.

## Acknowledgments

We thank Drs. Randy Haun and Ya-Wen Liu for their critical comments on this manuscript. We thank laboratory member Meng-Chen Tsai for preparing liposome floating assay and Dr. Kuan-Jung Chen for preparing phosphorylation reagents. This work was financially supported by grants from the Ministry of Science and Technology in Taiwan (ROC) [NSTC112-2320-B-002-019-MY3], National Health Research Institute, Taiwan (NHRI-EX113-11306BI) and the Center of Precision Medicine by the Ministry of Education in Taiwan to F.-J.S.L. We thank the Proteomics Core Laboratory at Chang Gung University for assistance with proteomic analysis.

## Data availability

The mass spectrometry data have been deposited to the ProteomeXchange Consortium via the PRIDE partner repository (Griss *et al*, 2016) with the data set identifier PXD054952 (reviewer account details: **Username:**; **Password:** PLQvzdltAjCe). All data generated or analyzed as part of this study are included in this published article and its supplementary information file. The mass spectrometry data (Dataset EV1) are provided with this paper.

---

**Fig. EV1.**
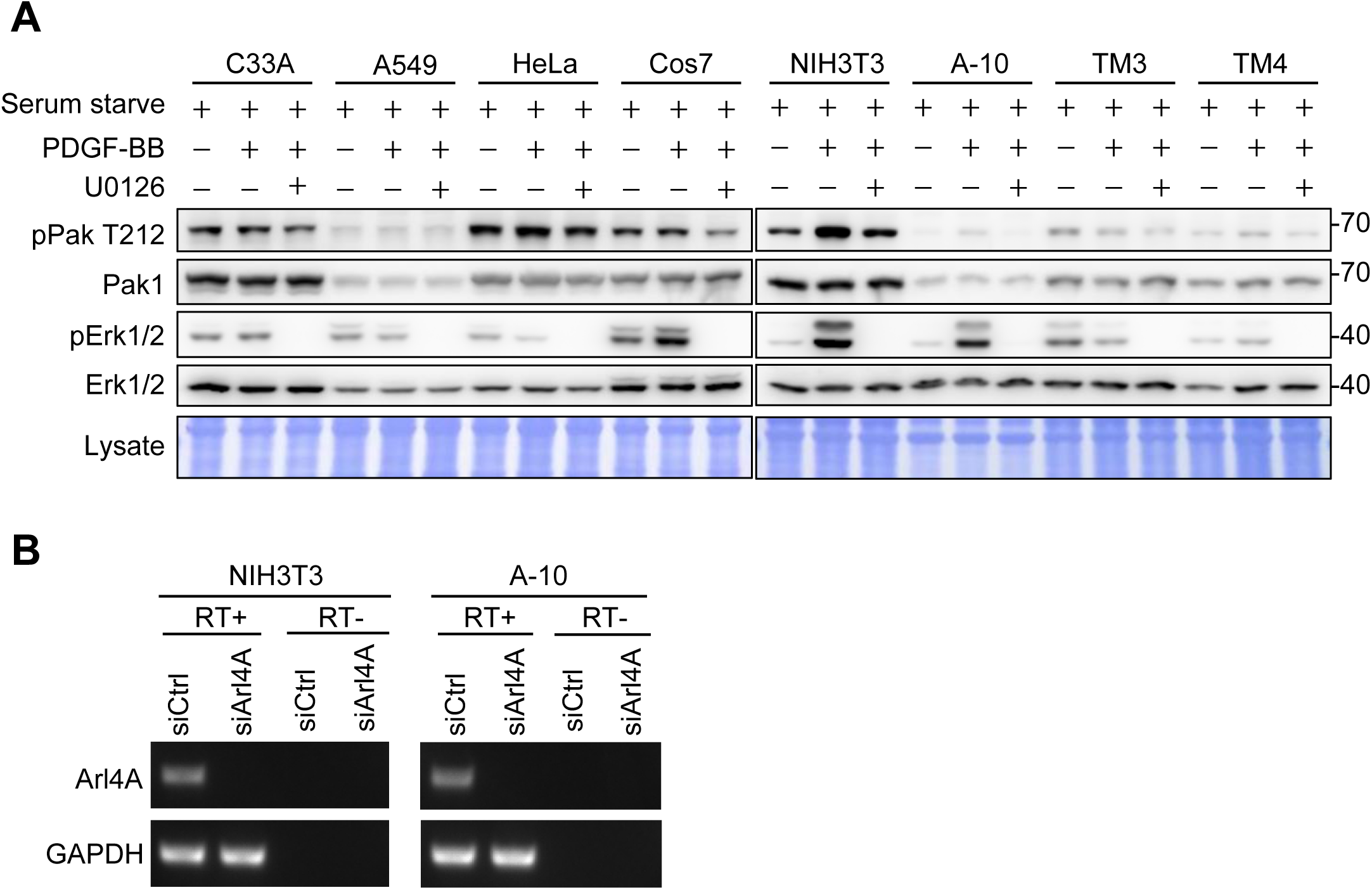
Screening of PDGF sensitive cell lines and confirmation of knockdown efficiency. (A) Indicated cell lines were serum staved and then treated with PDGF-BB at 20 ng/mL for 10 min or either co-treated with U0126 (10 uM). The indicated proteins were blotted with specific antibodies. Total cell lysates were stained with Coomassie blue. (B) NIH3T3 and A-10 cell lines knocked down with siCtrl and mouse siArl4A were serum staved and then treated with PDGF-BB (20 ng/mL) for 10 min. Total RNA was extracted from these cells and then subjected to RT-PCR. Using Arl4A-specific primers, the PCR products were analyzed on a 1.5% agarose EtBr gel for UV detection representing the mRNA levels of Arl4A.

**Fig. EV2.**
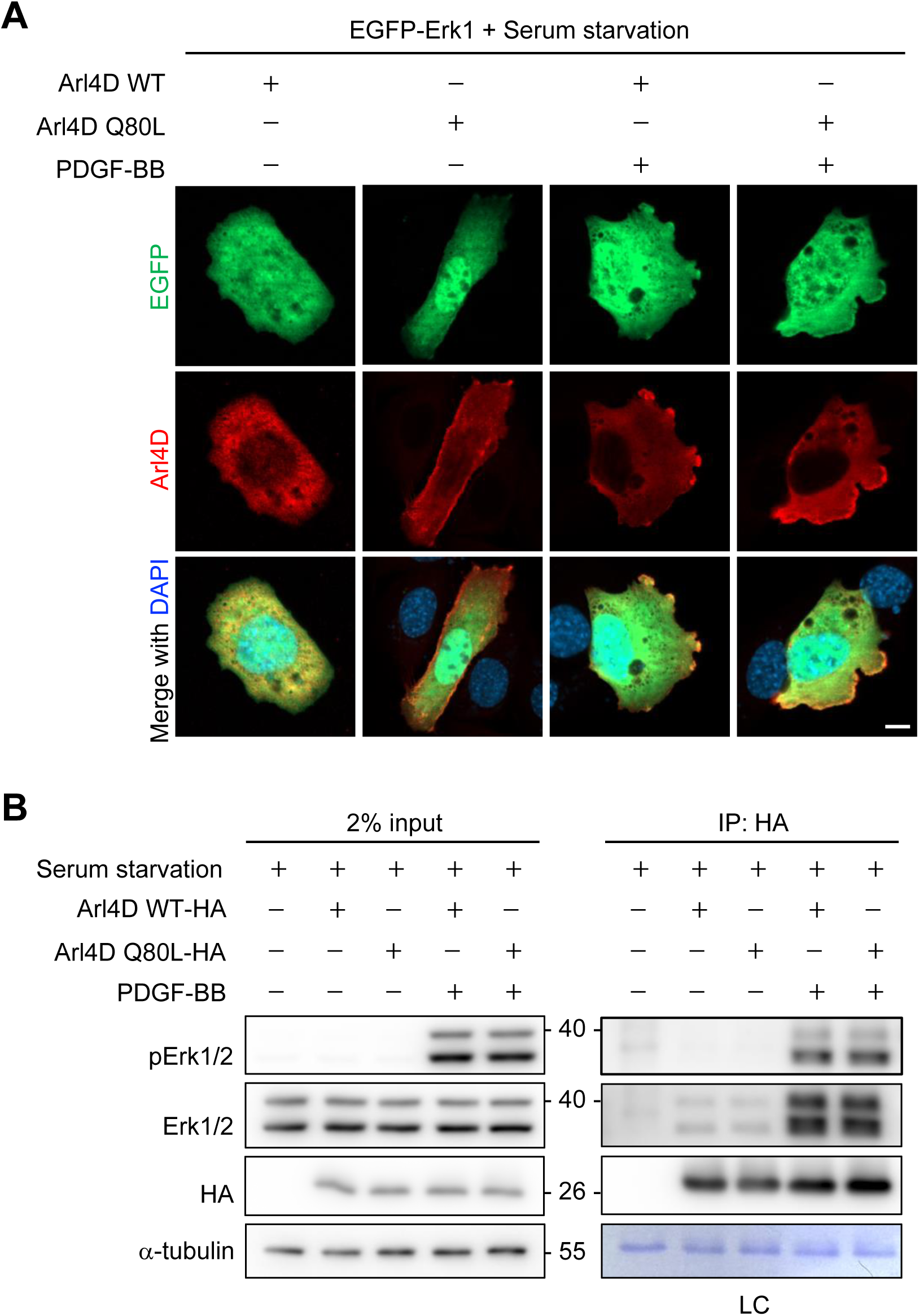
Arl4D interacts with activated Erk1/2 under PDGF signaling. (A) NIH3T3 cells transfected with indicated proteins were serum starved, treated with PDGF-BB at 20 ng/mL for 0 or 10 mins and stained with anti-Arl4D (red) antibodies and DAPI (blue; stains the nuclei). Scale bar, 10 mm. (B) NIH3T3 cells transfected with empty vector pSG5, Arl4D WT-HA or Arl4D Q80L-HA were serum starved, treated with PDGF-BB (20 ng/mL) for 10 min and was subjected to DSP-crosslinker-treated Co-IP.

**Fig. EV3.**
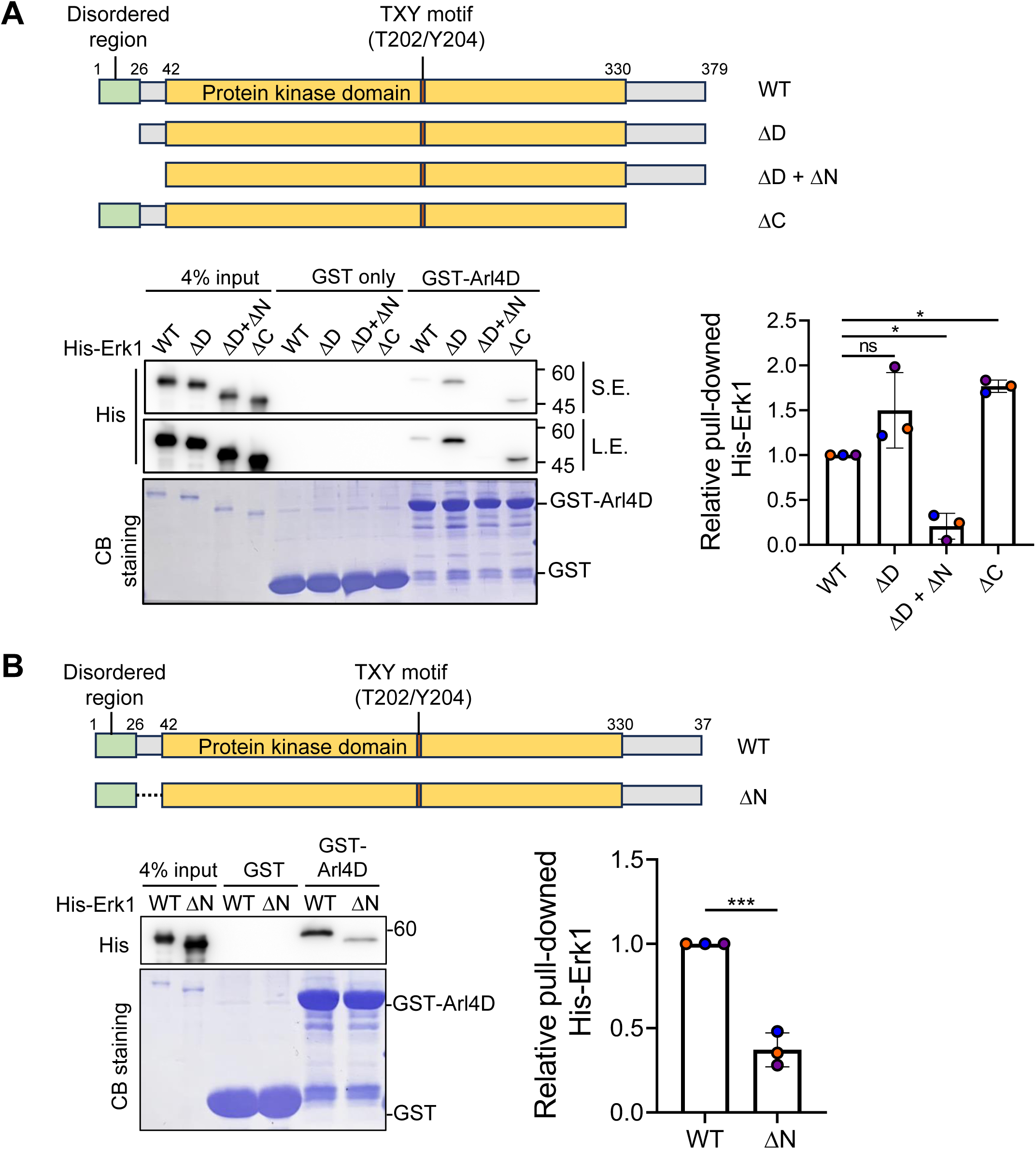
Identification of Arl4D-binding-defective Erk1/2. (A and B) Cartoon representations show the Erl1 truncations (ΔD: deletion of the disorder region; ΔN: deletion of the space between the disorder region and the kinase domain; ΔC: deletion of the region downstream of the kinase domain). In vitro binding of His-tagged Erk1 proteins with GST or GST-Arl4D. His-Erk1 proteins pulled down from GST fusion proteins were analyzed by Western blotting. Equal amounts of GST proteins were detected by staining with CB (Coomassie blue). The His signals of the pulled-down His-Erk1 proteins were quantified in the dot plots; the error bars indicate the mean ± (n=3). *P < 0.05; ***P < 0.001 (one-way ANOVA with Tukey’s post-hoc multiple comparison test in (A), two-tailed unpaired Student’s t-test in (B)).

**Fig. EV4.**
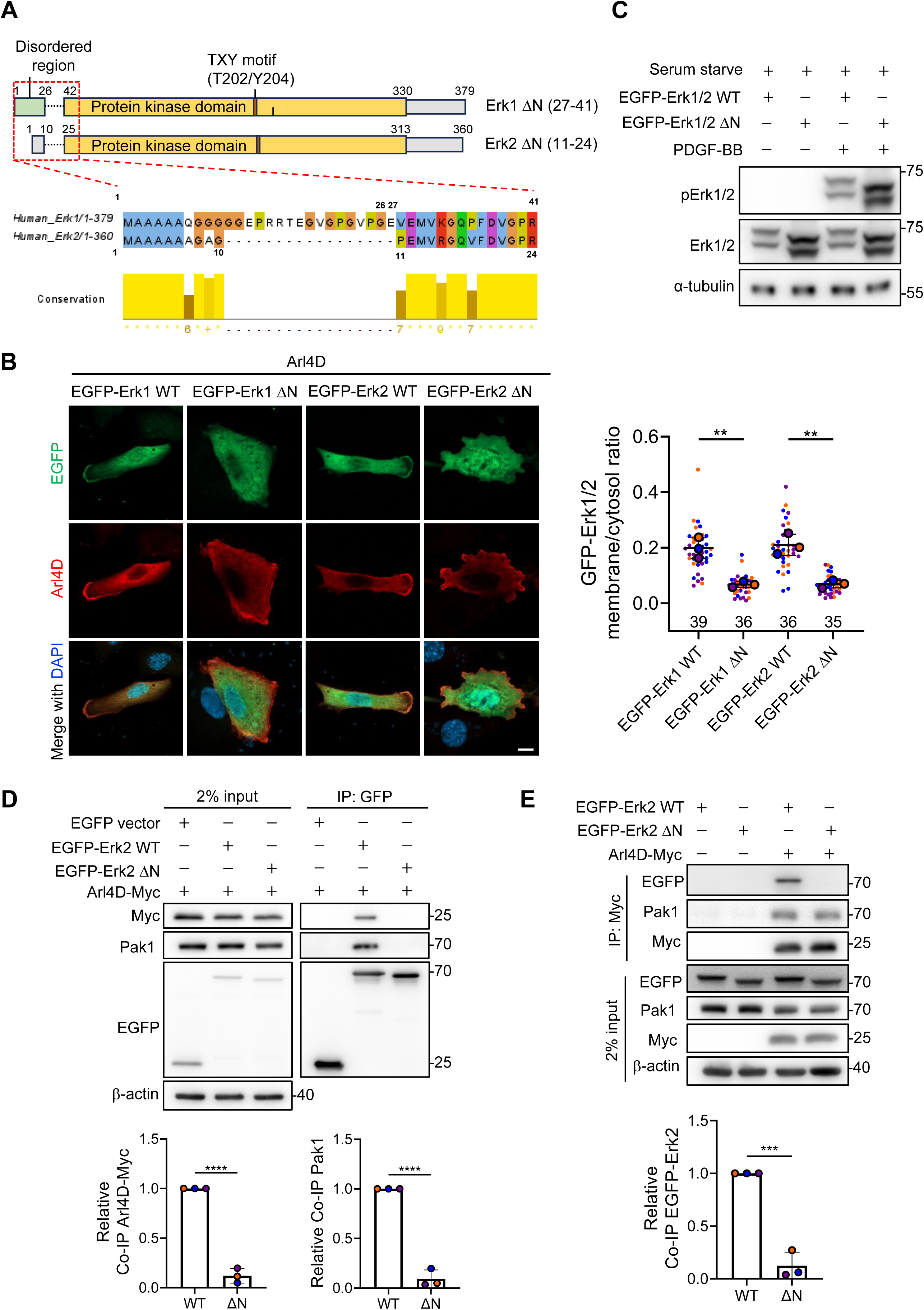
Erk1 and Erk2 share the same N-terminal region to interact with Arl4D. (A) Jalview software shows the alignment and conservation between Erk1 and Erk2. (B) NIH3T3 cells transfected with the indicated proteins were stained with anti-Arl4D antibodies (red) and DAPI (blue; stains the nuclei). Scale bar, 10 mm. The ratio between plasma membrane and cytosol of GFP-Erk1/2 in each group was quantified as described in Materials and Methods, and the results were shown in the dot plots with error bars indicating the mean ± SD (n=3, the cells analyzed in each biological replicate were marked in the same color, and the total number of cells is indicated in the graph). **P<0.01 (one-way ANOVA with Tukey’s post hoc multiple comparison test). (C) NIH3T3 cells transfected with EGFP-Erk1/2 WT or EGFP-Erk1/2 ΔN were serum starved and treated with PDGF-BB (20 ng/mL) for 10 min. Immediately harvested cell lysates were subjected to Western blotting for the indicated proteins. (D) NIH/3T3 cells transfected with pEGFP-C3, Arl4D-WT-Myc, EGFP-Erk2 WT, or EGFP-Erk2 ΔN were subjected to Co-IP. The Co-IP signals of endogenous Pak1 and Arl4D-Myc were normalized with GFP-Erk2 proteins after subtracting the background signal of the pEGFP-C3 group. (E) NIH/3T3 cells transfected with empty vector pSG5, Arl4D-Myc, EGFP-Erk2 WT or EGFP-Erk2 ΔN were subjected to Co-IP. The Co-IP signals of EGFP-Erk2 were normalized with Arl4D-WT-Myc proteins after subtracting the background signal of the empty vector control group. (D and E) The quantified results are shown in the dot plots, with the error bars indicating the mean ± SD (n=3). ***P<0.001; ****P < 0.0001 (two-tailed unpaired Student’s t-test).

**Fig. EV5.**
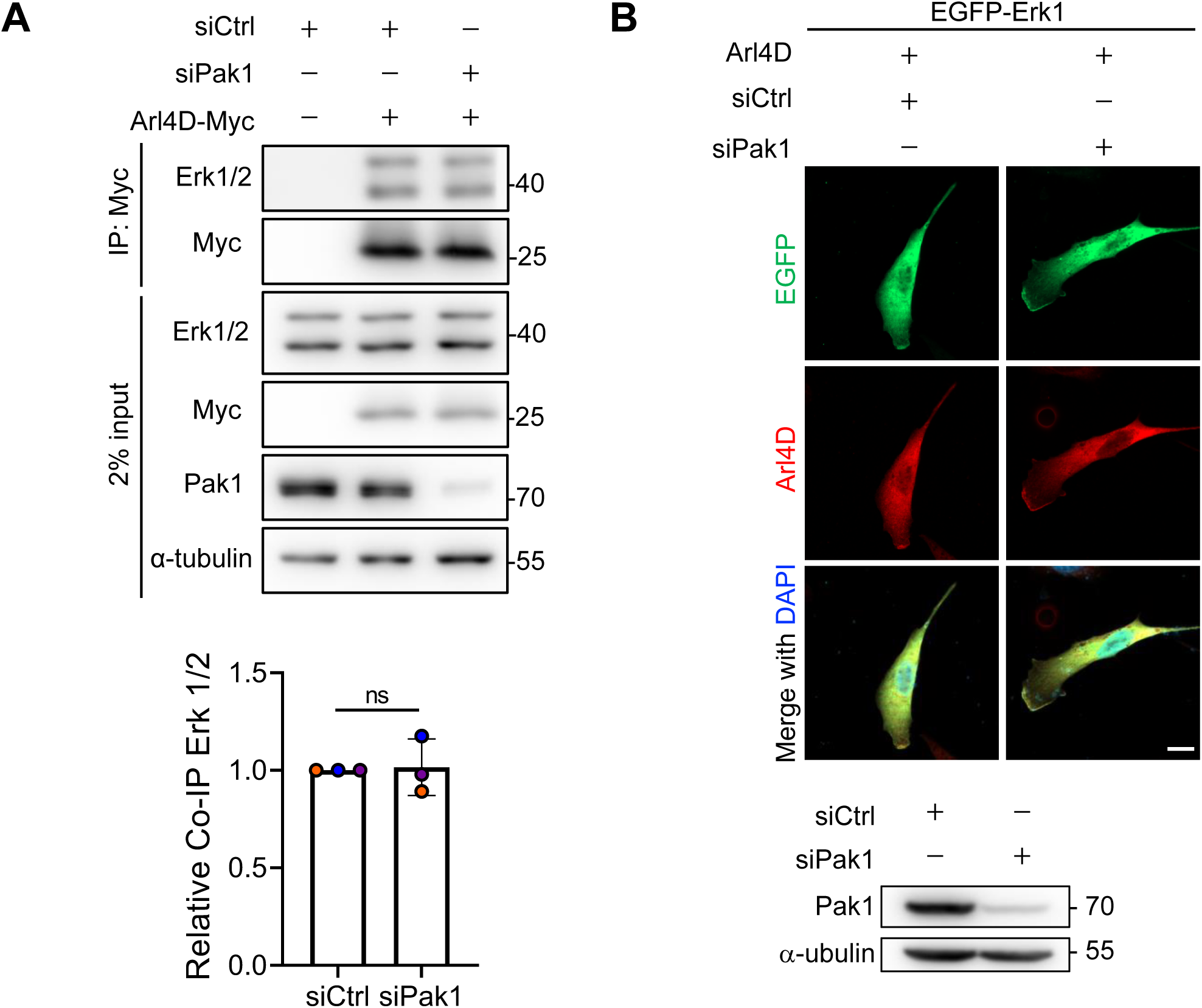
Arl4D interacting with Erk1/2 is independent of Pak1. (A) NIH3T3 cells knocked down with siCtrl or mouse siPak1 RNA and transfected with the empty vector pSG5 or Arl4D-Myc were subjected to Co-IP. The Co-IP signals of the endogenous Erk1/2 proteins were normalized with the Arl4D-Myc proteins after subtracting the background signal of the empty vector control group. The quantified results are shown in the dot plots with error bars indicating the mean ± SD (n=3, two-tailed unpaired Student’s t-test). (B) NIH/3T3 cells knocked down with siCtrl or mouse siPak1 and transfected with the indicated proteins were stained with anti-Arl4D antibodies (red) and DAPI (blue; stains the cell nuclei). Scale bar, 10 mm. The right panel shows the expression level of indicated protein.

**Fig. EV6.**
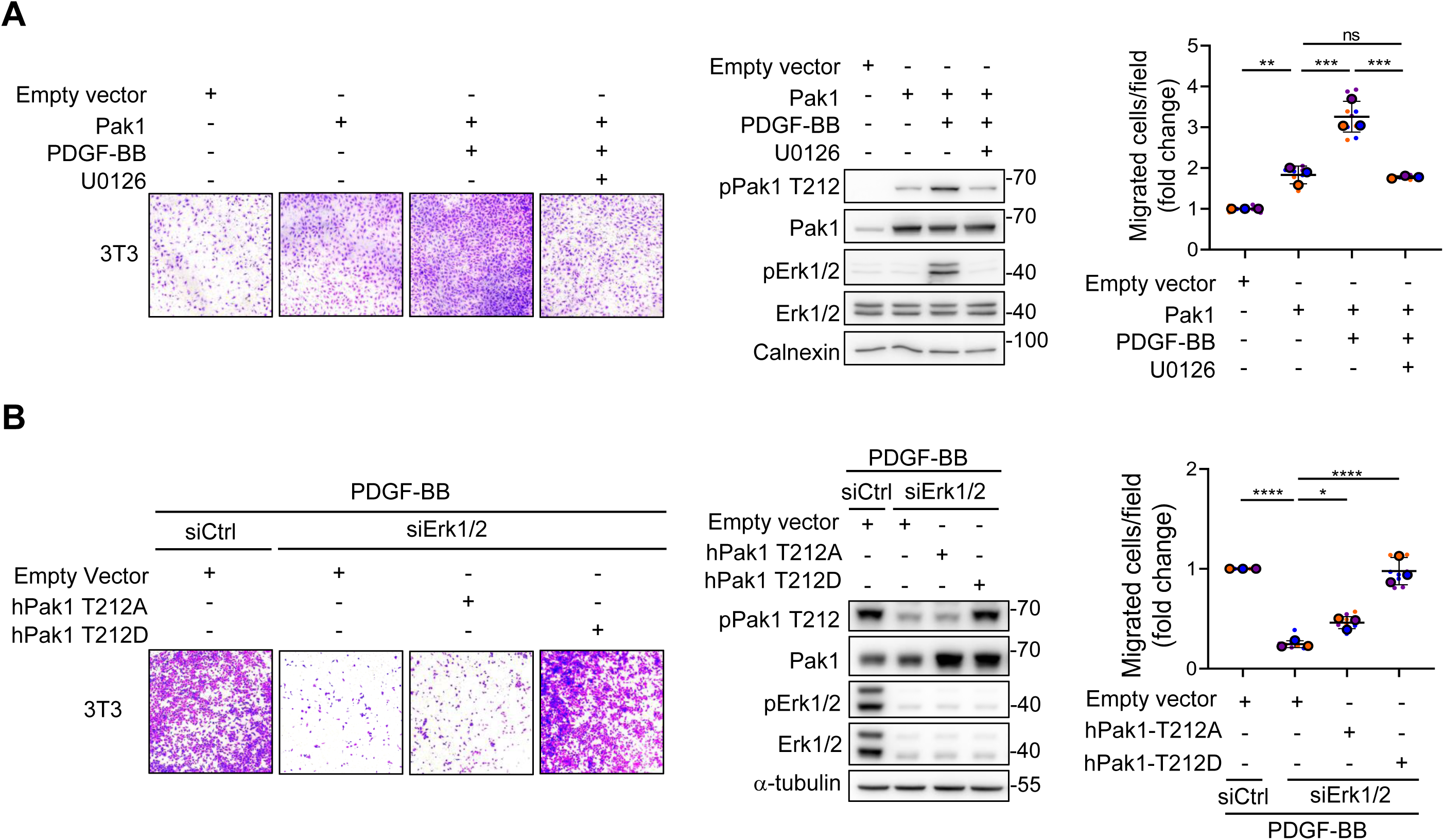
Pak1 T212 phosphorylation is important downstream signal under PDGF-Erk1/2 to promote cell migration. (A) NIH3T3 cells transfected with empty vector pcDNA3.0 or Pak1 were serum starved and treated with PDGF-BB (20 ng/ml) or co-treated with U0126 (10 uM). Western blotting detected the indicated proteins. (B) NIH3T3 were knocked down with siCtrl or mouse siErk1/2 RNA and transfected with empty vector pcDNA3.0, hPak1-T212A or hPak1-T212D. The cells were kept serum-free and treated with PDGF-BB (20 ng/ml). Western blotting detected the indicated proteins. (A and B) Cells migrated under the transwells were stained with crystal violet and photographed for 3 fields per biological replicate and calculated in dot plot with mean ± SD (n=3). *P<0.05; **P<0.01; ***P<0.005; ****P<0.001 (one-way ANOVA with Tukey’s post hoc multiple comparison test).

